# Outside-in progression of heterochromatin replication and exclusion of CDC45 from the PCH domain in *Drosophila*

**DOI:** 10.64898/2026.08.13.744572

**Authors:** Collin Hickmann, Srigokul Upadhyayula, Gary H. Karpen

## Abstract

Heterochromatin replication isn’t random or uniform, but occurs in a characteristic spatial and temporal pattern. Previous studies produced conflicting models for pericentric heterochromatin (PCH) replication, suggesting either that heterochromatic sequences translocate to the domain surface for replication, or that replication can also occur internally through localized decondensation. To distinguish true overlap from peripheral enrichment around an irregular PCH domain, we developed FOC-Map, a colocalization analysis approach that combines segmentation of one channel with binning of the other. Applying FOC-Map to three-dimensional Airyscan imaging of cultured *Drosophila* cells, we find that replication foci at the onset of late S-phase are confined to the outer boundary of the PCH domain, forming a shell-like pattern with little overlap into the HP1a-rich interior. As late S-phase progresses, replication foci are observed within the domain, localizing to low-HP1a regions interspersed between more condensed regions. We then assessed the distribution of CDC45, a rate-limiting replication initiation factor, and found that CDC45 foci are depleted from the PCH domain throughout the cell cycle. We propose that low levels of CDC45 within HP1a-rich PCH limit replication initiation to the domain periphery, giving rise to the shell-like pattern of replication foci that progressively works inward until PCH replication is complete.

## Introduction

Pericentric heterochromatin (PCH) is defined by enrichment for repeated DNAs, and the histone modification H3K9me2/3 and its reader protein HP1a, which together maintain a condensed, largely transcriptionally repressed chromatin state^1,2^. In *Drosophila melanogaster* and other metazoans, heterochromatic sequences coalesce into a membraneless biocondensate known as the PCH domain, formed through phase separation or similar biophysical mechanisms, and exhibiting liquid-like properties and selective permeability to macromolecules^3,4^. Despite its compaction, PCH must be faithfully replicated every cell cycle, and it does so with a characteristic delay relative to euchromatin, a conserved feature of metazoan cell biology whose mechanistic basis remains incompletely understood. HP1a plays a central role in this delay: targeting HP1a to sequences that are not ordinarily heterochromatinized is sufficient to delay their replication, and its loss reduces the extent of late replication beyond what can be explained by other known regulators such as RIF1^5,6^. How HP1a imposes this delay, whether the physical properties of the PCH domain it organizes contribute to it, and how the mechanisms responsible for the temporal dynamics of PCH replication relate to its spatial organization are open questions.

Previous studies in mammalian cells have produced two partially overlapping models describing where heterochromatin replication occurs relative to the PCH domain. Foundational work in fixed mammalian cells observed that sites of DNA replication (replication foci) during late S-phase are confined to the surface of the PCH domain, and suggested that that heterochromatic sequences may need to relocate to the domain surface in order to be replicated^7^. Subsequent studies using higher-resolution imaging found that replication foci also appeared within less compacted regions inside the domain^8^. A later study in live cells described a temporal progression in which replication begins at the periphery and later extends into the interior within locally decondensed channels, without evidence for large-scale translocation^9^.

Heterochromatin replication dynamics have also been characterized in the accelerated cell cycles of early *Drosophila* embryos, where replication was observed in decondensed regions adjacent to the PCH-domain^10^. However, it remains unclear how these findings relate to the models developed in mammalian cells and whether they hold true outside of the rapid embryonic cell cycles.

Prior efforts to characterize the spatial organization of PCH replication have relied predominantly on markers of replication elongation, such as PCNA and nucleotide analogs, to visualize sites of ongoing DNA synthesis. While informative for characterizing where replication is actively occurring, these approaches provide limited insight into the role of replication initiation in shaping these dynamics. The comparatively few studies that have addressed initiation in this context have largely focused on regulators that modulate the expression or activity of initiation factors, such as the role of RIF1 in antagonizing DDK-dependent origin activation^5,11^. However, few have directly examined how the spatial distribution of the initiation factors themselves might influence the dynamics of PCH replication. The condensate-like properties of the PCH domain have been shown to influence the distribution of macromolecules within it through selective enrichment and exclusion^4,12^. Whether replication initiation factors are also affected by these properties, and whether their distribution with respect to the PCH domain influences the dynamics of heterochromatin replication, has not been directly examined. To begin addressing these questions, we focused on CDC45, a conserved component of the CMG helicase complex that is assembled at licensed origins to initiate DNA synthesis^13,14^. CDC45 is present in limiting amounts relative to the MCM2-7 helicase loaded at origins, and its availability is a primary constraint on the rate of origin firing^15–18^, making its distribution within the nucleus particularly likely to influence local rates of replication initiation, and by extension, the dynamics of PCH replication.

Here, we set out to characterize the spatial dynamics of PCH replication in cultured *Drosophila* S2R+ cells and to investigate the distribution of CDC45 relative to the PCH domain. Using three-dimensional time-lapse Airyscan microscopy, we tracked the progression of replication foci through late S-phase. Quantification of the spatial relationship between replication factors and HP1a in live studies is challenging using existing analysis methods due to the dynamic behaviors of the PCH condensate. Therefore, we developed FOC-Map, a new method for the quantitation and visualization of colocalization, exclusion, and proximity of objects in one channel with signal in the other. Our analysis demonstrated that replication foci are initially concentrated at the periphery of the PCH domain at the onset of late S-phase, forming a characteristic shell-like pattern, and that replication later extends into low-HP1a regions interspersed throughout the increasingly fragmented domain. We then assessed the distribution of CDC45 by immunofluorescence and found that CDC45 foci are depleted within the PCH domain throughout the cell cycle. Together, these results suggest a model in which the reduced availability of CDC45 and potentially other initiation factors within the HP1a-rich PCH domain limits the capacity for replication initiation in the domain interior, giving rise to the peripheral pattern of replication observed at the onset of late S-phase and contributing to the progressive, ‘outside-in’ replication of heterochromatin.

## Results

### FOC-Map: a method for quantifying and visualizing colocalization, exclusion, and proximity in microscopy images

To determine whether heterochromatin replication occurs primarily at the periphery of the PCH domain or also in its interior, we used Airyscan microscopy to perform three-dimensional time-lapse imaging of replication foci (visualized with mScarletI-PCNA) and the PCH domain (visualized with mEGFP-HP1a) in cultured *Drosophila* S2R+ cells progressing from early through late S-phase (Fig. 2A). Analysis of these images first required a method for quantifying the enrichment of features in one channel adjacent to those in the other. However, existing colocalization methods are sensitive only to direct co-occurrence, as in object-based approaches like the Manders Colocalization Coefficient (MCC), or correlation between channels, as in pixel-based approaches like the Pearson Correlation Coefficient (PCC)^19–21^. The diffuse boundaries and heterogeneous internal intensity of the PCH domain further complicate the application of approaches like MCC that require segmentation of both channels, and while line intensity profiles can reveal peripheral relationships qualitatively, they are not volumetric and cannot be scaled to larger datasets^22^.

To address these limitations, we developed FOC-Map (Fractional Overlap Coefficient Map), an approach that captures not only direct overlap and exclusion but also enrichment of one signal adjacent to the other (Fig. 1A). Unlike other object-based measures of colocalization that require segmentation for both channels in an image, FOC-Map segments structures in only one channel (e.g., replication foci) and quantifies their distribution across percentile intensity bins in the other channel (e.g. HP1a/PCH). Quantifying the normalized overlap of the segments with each percentile intensity bin makes it possible to determine how objects in one channel are distributed with respect to regions of varying intensity in the other, while avoiding the need for a single arbitrary threshold. Voxels in the region of interest (i.e. the nucleus) are also binned by their proximity to the voxels of each intensity bin, producing a normalized distance profile that shows when segments in one channel are enriched in close proximity to high-intensity regions of the other.

**Figure 1.**
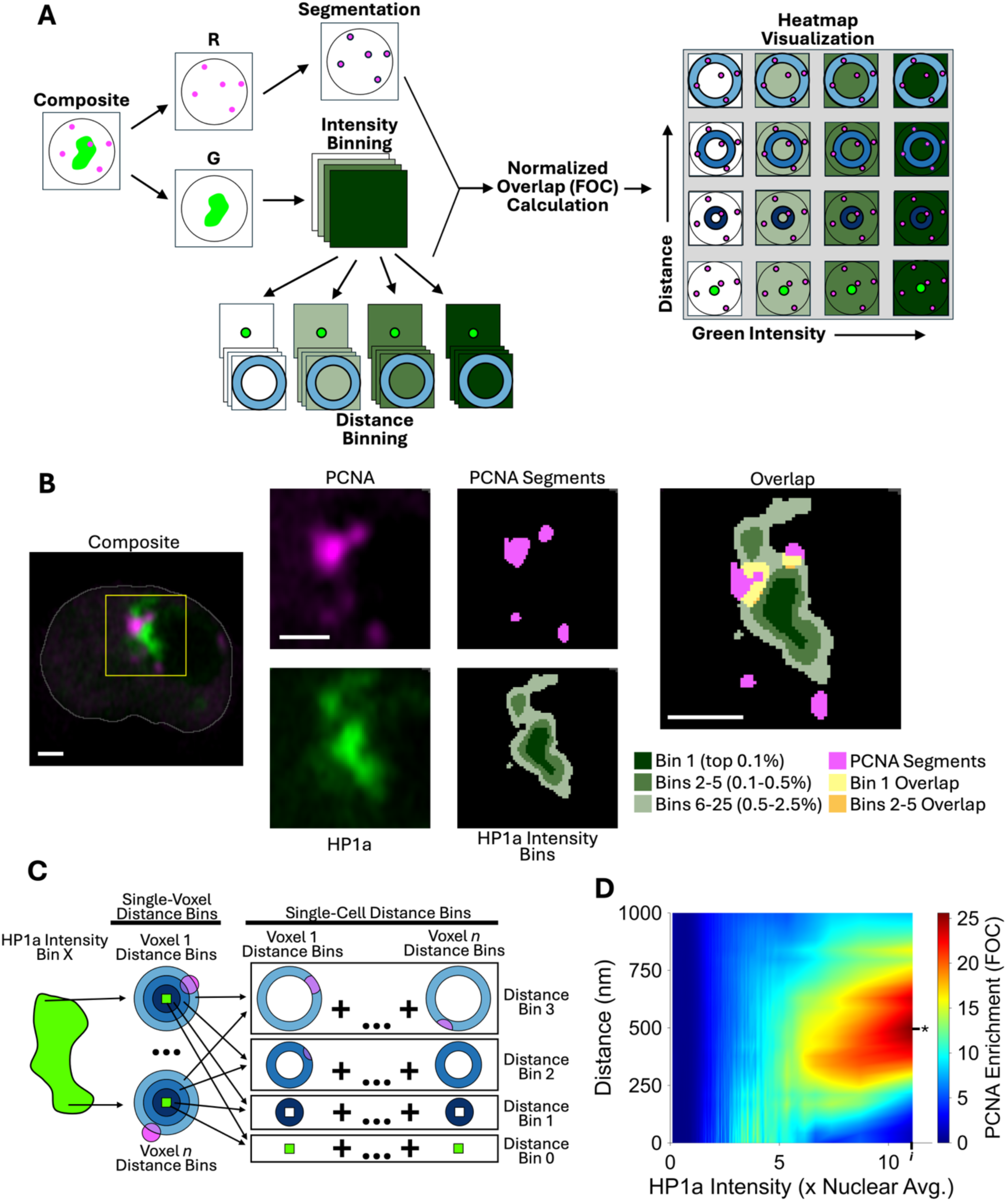
Schematic overview of FOC-Map and visualization of output in a single cell exhibiting peripheral localization of replication foci. **A.** Simplified schematic showing the workflow of FOC-Map. While features in the red channel (e.g. replication foci) are segmented, the green channel (e.g. PCH) undergoes intensity binning, and a series of distance bins is made for each intensity bin using the distance between voxels in each intensity bin of the green channel and those in segments of the red channel. The FOC is used to measure the normalized overlap between the red segments and each of the green bins. These FOC values are then displayed in a heatmap. While the bin FOC values are represented graphically here, their values are conveyed by color-scaling in actual heatmaps, as shown in subsequent figures. **B.** PCNA segments and HP1a Percentile Intensity Bins visualized in a single Z-plane of a live S2R+ *Drosophila* cell in late S-Phase. Replication foci (magenta, visualized with mScI-PCNA) border the PCH domain (green, visualized with mEGFP-HP1a) but lack extensive overlap in the high-intensity core of the domain. Intensity bins are grouped for visibility and conceptual clarity; scale bars are 1 μm. The nucleus is outlined in grey in the first panel, while the yellow inset is magnified and displayed in all subsequent panels. **C.** Graphical representation of the distance binning procedure for a single intensity bin. Concentric distance bins are generated from each voxel within the selected intensity bin and the distance bins from each voxel are then pooled by their radii (distance from central voxel). In practice, the full set of distance bins for each voxel spans the entirety of the ROI (e.g. the nucleus) such that each voxel of the ROI is binned separately based on its distance to each voxel in the intensity bin. This process is then repeated for each intensity bin. **D.** The resulting FOC-Map for the cell shown in B. The asterisk indicates the region of peak association with PCNA segments (highest FOC value) ∼500 nm away from the top intensity bin, while the “*i*” tick at the right side of the x-axis shows the highest HP1a intensity bin, where the FOC has a value of 2.5.

At the core of FOC-Map is the Fractional Overlap Coefficient (FOC), a normalized measure of overlap between the segments in one channel and each bin in the other. Because the degree of overlap expected by chance depends on the volumes of the segments, the bins, and the region of interest containing them, the FOC normalizes the observed overlap against that expected under a null model of random distribution, producing a measure that reflects spatial association independent of these volumes (see Methods). If no spatial relationship exists between features in the two channels and their distribution is random with respect to each other, the fraction of red volume overlapping green will, on average, be equal to the fraction of the ROI overlapping green, producing an FOC of 1. An FOC of 2 would indicate the fractional overlap of the bin is twice that of the broader ROI, and twice that which would be expected in the case of random distribution. By accounting for differences in both bin and segment volume, the FOC provides an intuitive measure of normalized overlap that allows meaningful comparisons across bins of variable size.

Percentile intensity binning groups the voxels within each nucleus by their relative HP1a intensity, bypassing the need for segmentation of the PCH domain and making it possible to measure how PCNA foci are distributed between regions of varying HP1a intensity (Fig. 1B). In addition to direct overlap, FOC-Map quantifies the enrichment of segments at varying distances from the voxels in each intensity bin.

Nuclear voxels are binned by their distance from the voxel with the greatest HP1a intensity, effectively creating a series of spherical shells radiating outward from this maximum intensity voxel (Fig. 1C). This process is repeated for each of the voxels in the highest intensity bin, and the resulting distance bins from each voxel in this intensity bin are pooled together. This is then repeated for each intensity bin, and the FOC is calculated for each combination of intensity bin and distance bin. Since binning is performed in two dimensions (intensity and distance), the results are plotted on a heatmap, with HP1a intensity bins along the X-axis and distance from the voxels of each intensity bin along the Y-axis, with the FOC for each bin encoded as color (Fig. 1D). The progression of colors from the bottom to the top of the heatmap at the far right displays how the FOC varies with distance from the voxels in the highest HP1a intensity bin.

To illustrate how these spatial relationships are represented in FOC-Map, we applied it to a cell (from the live-cell time-lapse dataset described below) in which replication foci are clustered at the periphery of the PCH domain with relatively little overlap in its high-HP1a core (Fig. 1B, D). The enrichment of replication foci at the periphery of the PCH domain is visible as a prominent peak in FOC values approximately 500 nm from the voxels of the highest HP1a intensity bin (Fig. 1D, asterisk), reaching a value of 25. The FOC at the highest intensity bin itself is roughly 2.5 (Fig. 1D, *i*), which is above the baseline of 1, but well below the peak in adjacent regions and lower intensity bins. Because both signals are confined to a small fraction of the nuclear volume, even modest overlap can produce FOC values above 1, making comparisons across the full set of bins more informative than any single value in isolation. For comparison, when FOC-Map is applied to a positive control for colocalization in which segmented PCNA foci are measured against PCNA intensity bins, the FOC reaches values over 100 in the brightest bins (Supplementary Fig. 1A). Conversely, in an early S-phase cell where replication foci are euchromatic and largely excluded from the PCH domain, the FOC in high-HP1a bins falls well below 1 (Supplementary Fig. 1B). Together, these examples illustrate that FOC-Map can distinguish colocalization, exclusion, and peripheral association, and that the spatial context provided by the full heatmap reveals relationships that would be difficult to capture with a single summary statistic.

### Late-S replication foci are first enriched at the periphery of the HP1a-rich PCH domain and later in interspersed low-intensity regions

Having established FOC-Map as a method for quantifying colocalization, exclusion, and peripheral association, we next applied it to the full live-cell time-lapse dataset to determine how the distribution of replication foci relative to the PCH domain changes over the course of late S-phase. Cells in early S-phase exhibited numerous small, widely distributed replication foci (Fig. 2A, −20% late-S progression), reflecting widespread euchromatin replication^23,24^. At the onset of late S-phase, replication foci underwent a pronounced spatial reorganization to form a distinctive shell-like pattern at the boundary of the HP1a-rich PCH domain (Fig. 2A, 20% late-S progression). As late S-phase progressed, the morphology of the PCH domain became increasingly fragmented and irregular, and replication began to occur at internal, low-intensity channels separating highly condensed HP1a-rich regions. To allow comparison between cells despite variation in late-S duration, we aligned the time-lapse images from each cell by their relative progression through late-S, expressed as a percentage. Percentile intensity binning was performed as before, but the HP1a-intensity bins were pooled between the cells of each timepoint and the FOC was calculated for each of the pooled bins, allowing a representative result to be determined for each timepoint while minimizing the contribution of noise in single cells.

**Figure 2.**
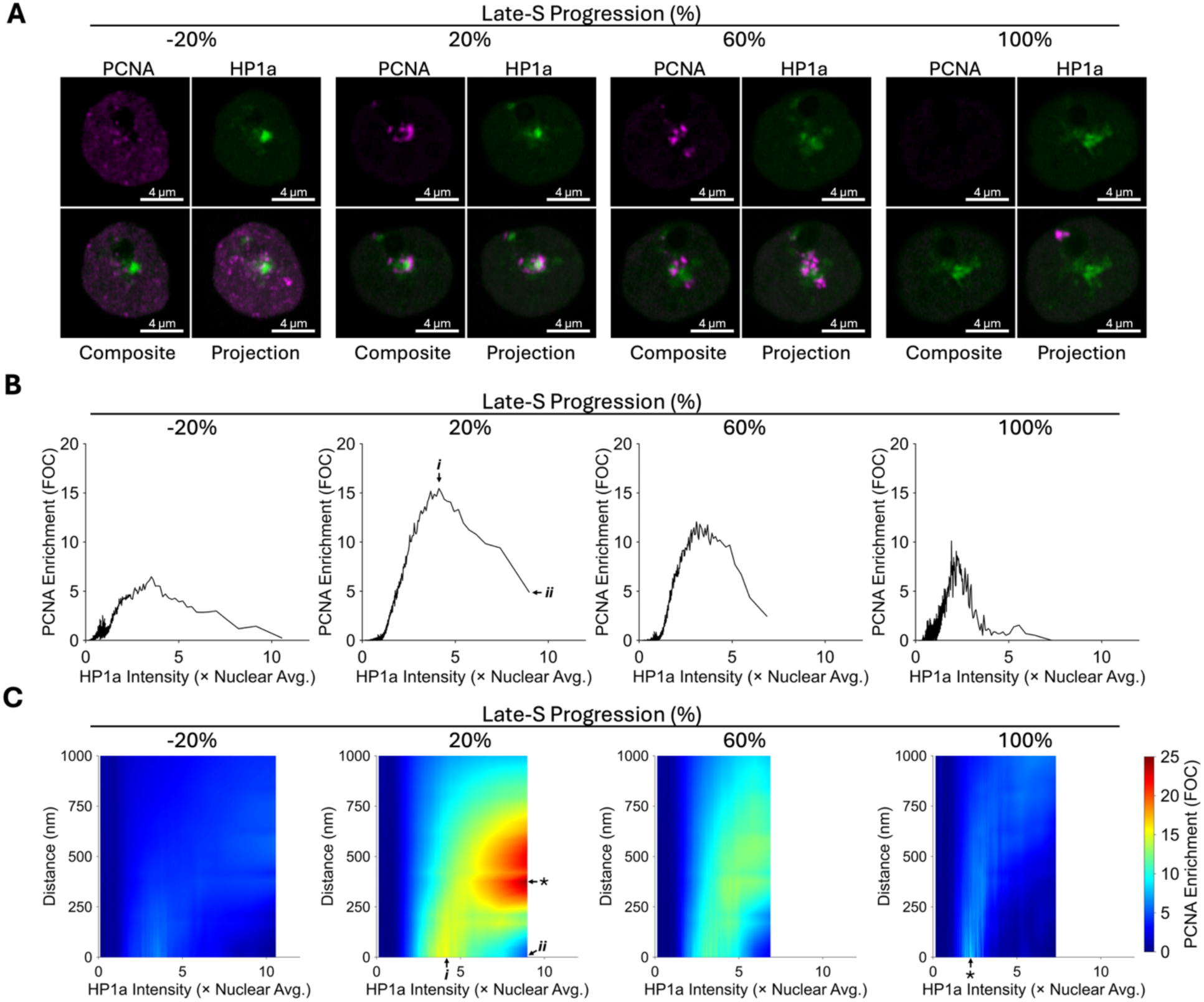
Late-S replication foci are enriched at the periphery of the PCH domain and in regions with relatively low levels of HP1a. **A.** Representative images from time-lapse analysis of cultured *Drosophila* S2R+ cells transiently expressing mScarletI-PCNA and mEGFP-HP1a, at each of the S-phase timepoints used in subsequent analysis. Replication foci form a shell at the periphery of the PCH domain at the onset of late-S, but occupy lower intensity regions interspersed between areas of greater intensity later in S-phase. Colocalization with the high-HP1a core of the PCH domain appears limited at all timepoints. Time lapses were temporally aligned by their relative progression through late S-phase, with the initial convergence of replication foci at the PCH domain defining the onset of late S-phase (0% late-S progression) and the last timepoint with replication foci defining the end of S-phase (100% progression). **B.** Pooled plots of fractional PCNA overlap vs. HP1a intensity at each timepoint (n = 10 nuclei). Nuclear voxels were binned into percentile groups with a bin size of 0.1%, the Fractional Overlap Coefficient (FOC) with PCNA was calculated for each, and the values were plotted against the average relative HP1a intensity of each bin. PCNA overlap with the high-HP1a bins increases dramatically at the 20% timepoint following the onset of late-S (arrow *ii*), but remains well below the peak overlap in much lower intensity bins (arrow *i*). **C.** Pooled FOC-Maps of fractional PCNA overlap (color) vs. HP1a percentile intensity bins (x-axis, 0.1% bin size) and binned distance from each intensity bin (y-axis, 43 nm bin size).

In early/mid S-phase (Fig. 2B, -20% late-S progression), PCNA foci were distributed across a broad range of intensity bins with relatively low HP1a intensity, resulting in a lower peak FOC compared to later timepoints, and consistent with the dispersed replication of euchromatin during this part of S-phase. At the onset of late S-phase, replication foci became strongly enriched in bins with less than half the average HP1a intensity of the top bin. The FOC in these moderate intensity bins reached a peak of 15 (Fig. 2B, 20% late-S progression, arrow ***i***), which is three times greater than the amount detected in the highest intensity bin (arrow ***ii***) and 15 times greater than would be expected if replication foci were randomly distributed throughout the nucleus. This enrichment in moderate intensity bins is consistent with the preferential localization of replication foci along the periphery of the PCH domain, where HP1a intensity is lower than in the domain’s core. The preferential association with lower intensity bins also holds true for later stages of late S-phase, although with lower peak FOC values that occur in bins with even lower relative HP1a intensity, consistent with the qualitative association with low intensity regions (Fig. 2B, 60 – 100% late-S progression).

We next sought to quantify how fractional PCNA overlap varied with distance from high-HP1a regions. Because PCNA foci appeared to form a shell surrounding the high intensity core of the PCH domain at the onset of late-S, we reasoned that the pattern of overlap observed in the intensity plots could be an indirect reflection of this spatial relationship. To test this, we applied the distance binning component of FOC-Map as described in the single-cell examples (Fig. 1C), but pooling across cells at each timepoint as was done for the intensity analysis. The resulting FOC-Maps display how the enrichment of PCNA foci varies with both HP1a intensity and distance from each intensity bin (Fig. 2C).

The pooled FOC-Map for the first timepoint (Fig. 2C, -20% late-S progression) shows a very broad distribution of replication foci without strong enrichment at any particular intensity class or distance, reflecting the dispersed replication of euchromatin throughout the nucleus in early/mid S-phase. In contrast, the FOC-Map for cells just after the onset of late S-phase shows a pronounced peak in FOC values 300 – 600 nm from the highest HP1a intensity bin (Fig. 2C, 20% late-S progression, asterisk), with nearly twice the FOC value of any intensity bin at zero distance. This indicates that PCNA is more highly enriched in the region adjacent to high-intensity HP1a voxels than at any particular intensity level, and reflects the “shell” configuration of PCNA foci along the boundary of the PCH domain visible in the images from this timepoint. The relationship between HP1a and distance provides further support for this conclusion.

While the distance to the FOC peak for the highest intensity bin is nearly half a micron, this distance decreases with HP1a intensity in lower bins, ultimately reaching zero for the intensity bins with the highest direct overlap, which have less than half the HP1a intensity of the top bins (Fig. 2C, 20% late-S progression, arrow ***i***). The association of PCNA foci with lower intensity bins at this time may be a consequence of the fact that the HP1a-rich core of the PCH domain is surrounded by voxels with relatively lower HP1a intensity that are still well above the nuclear background. This shows that at the onset of late S-phase, replication foci directly overlap regions of low to moderate HP1a intensity bordering higher intensity regions.

At later timepoints, the enrichment of replication foci adjacent to high intensity HP1a regions decreases, and the trend in the FOC-Map is dominated by the association of replication foci with lower intensity classes, rather than their proximity to high intensity regions (Fig. 2C, 100% late-S progression, asterisk). This is consistent with the observed change in the distribution of replication foci in the latter half of S-phase, where foci are primarily observed in regions of low HP1a intensity interspersed between higher intensity regions.

Rather than simply measuring the amount of overlap between replication foci and the PCH domain, FOC-Map made it possible to visualize the aggregate distribution of replication foci with respect to regions of varying HP1a intensity. This revealed that PCNA foci are most strongly enriched not within high-HP1a regions themselves, but in close proximity to them, concentrated in regions with less than half the peak HP1a intensity, at distances of 300 - 600 nm from the highest intensity voxels. This enrichment was strongest shortly after the onset of late S-phase, when the qualitative shell-like pattern was most apparent, and diminished at later timepoints as replication foci became increasingly associated with low-HP1a channels interspersed throughout the fragmenting domain. Throughout this progression, direct overlap with the HP1a-rich core remained consistently low relative to the enrichment observed in adjacent and lower-intensity regions. These patterns are broadly consistent with qualitative observations made here and in prior work, but the quantitative detail provided by FOC-Map goes beyond what could be determined by visual inspection alone. In particular, the ability to distinguish peripheral enrichment from direct overlap and to track how this relationship changes with both intensity and distance reveals spatial structure in the data that would otherwise be obscured. The near-absence of replication foci from the domain interior at the onset of late S-phase raises the question of what constrains replication to the periphery during this initial phase, and how this pattern might relate to events upstream of the replication elongation visualized as replication foci.

### CDC45 foci are depleted within the PCH domain

We reasoned that the observed restriction of replication to the surface of the PCH domain at the onset of late S-phase could result from decreased replication initiation in its interior. While there is no established method for the visualization of initiation sites *in situ*, the initiation factor CDC45 is known to be rate limiting for replication initiation, and its distribution within the nucleus would therefore be expected to influence local rates of initiation. Motivated by the past work showing that the PCH domain may partially exclude some factors by selective permeability^4^, we hypothesized that CDC45 is excluded from the PCH domain. To test this hypothesis, the distribution of CDC45 and its relationship to the PCH domain was characterized by immunofluorescence staining in fixed S2R+ *Drosophila* cells. Immunofluorescence allowed endogenous CDC45 to be visualized without the artefacts that are known to occur with its ectopic expression in live cells^16^. Cell cycle stage was determined using a combination of EdU-labelled replication foci and DNA content (integrated nuclear DAPI), allowing cells in G1 phase, early S-phase, late S-phase, and G2 phase to be differentiated from each other.

Staining for CDC45 revealed a punctate nuclear distribution in all stages of the cell cycle, but with highly variable intensity between cells (Fig. 3A). Although prior studies have shown that CDC45 remains associated with the replisome during elongation^25^, CDC45 puncta were distinct from replication foci visualized by EdU labelling (Fig. 3A), with minimal apparent overlap between the two signals. It is unclear whether the apparent lack of visible CDC45 enrichment at elongating replication foci indicates that only a small fraction of CDC45 is associated with elongating replication forks at any given time in S-phase, or instead suggests that our antibody has difficulty accessing or recognizing the epitope in elongation-associated CDC45. Since the current aim is focused solely on CDC45’s role in replication initiation, the negligible level of signal from elongation-associated CDC45 simplifies the quantitation and interpretation of the results.

**Figure 3.**
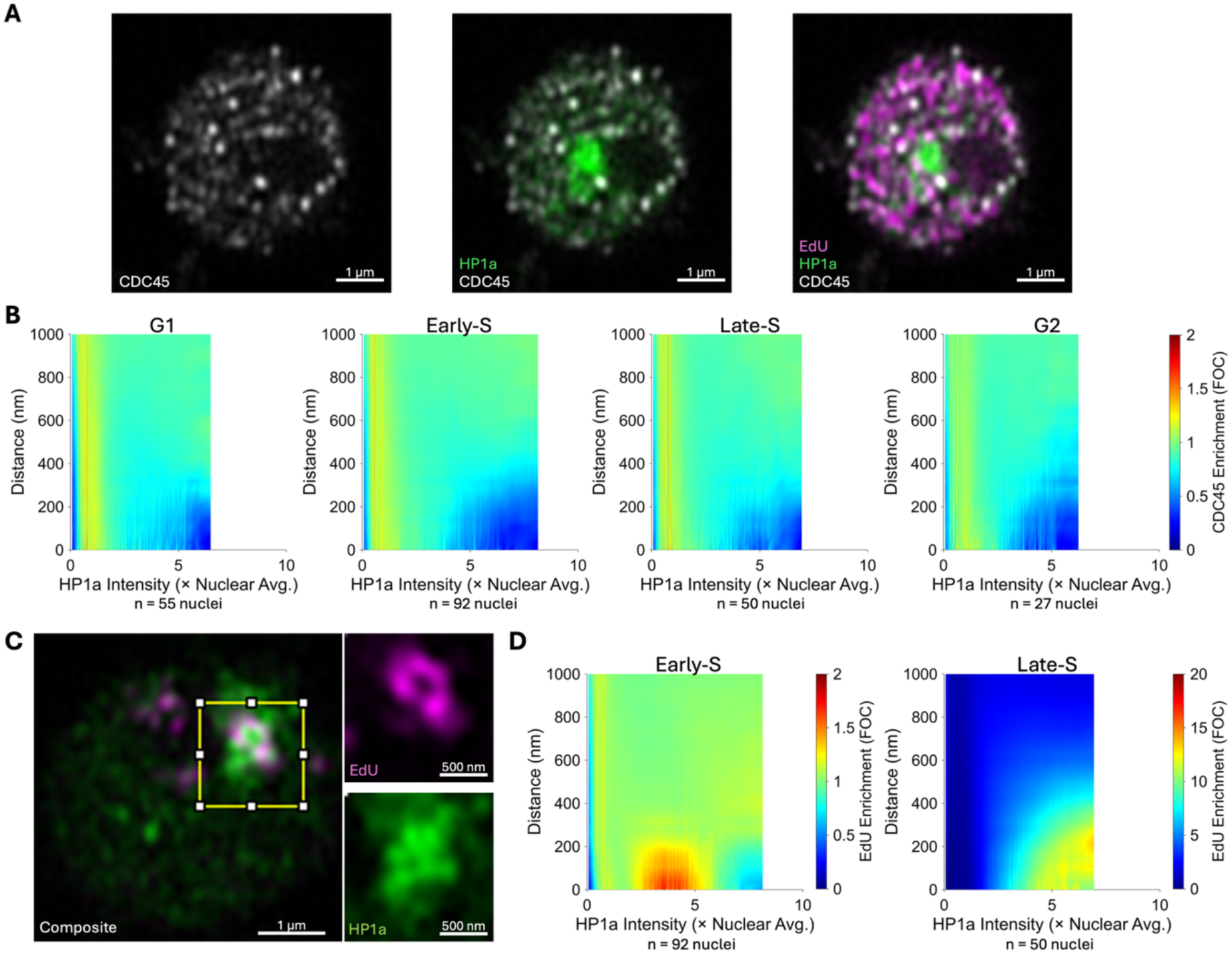
CDC45 foci are excluded from the PCH domain. **A.** Representative immunofluorescence image of a fixed S2R+ cell in early S-phase stained for HP1a (green) and CDC45 (gray) after pulse-labelling with EdU to mark replication foci (magenta). CDC45 forms nuclear foci that seem to be present at reduced levels within the PCH domain relative to the rest of the nucleus. **B.** Pooled FOC-Map of Fractional CDC45 overlap with nuclear voxels binned by percentile HP1a intensity (X-axis, 0.05% bin size) and distance from each intensity bin (Y-axis, 35.5 nm bin size). During all phases of the cell cycles, overlap of CDC45 foci with high HP1a-intensity bins is strongly reduced below the nuclear average of 1.0. **C.** Representative image of EdU foci and HP1a in late S-Phase. EdU foci appear to form a shell around high-HP1a regions that is qualitatively similar to PCNA in live cells, but the reduced scale of the structures in fixed cells approaches the resolution limit and increases the apparent overlap. **D.** Pooled FOC-Maps displaying the normalized overlap (FOC) of EdU foci with nuclear voxels binned by percentile HP1a intensity (X-axis, 0.05% bin size) and distance from each intensity bin (Y-axis, 35.5 nm bin size) in the same cells quantified in Fig. 3B.

Segmentation of CDC45 foci allowed colocalization with HP1a to be measured by the same FOC-Map approach used for PCNA, while also reducing the contribution of low intensity background and autofluorescence. Quantitation revealed that fractional overlap of CDC45 with high-HP1a regions is extremely low. The FOC of the highest HP1a intensity bin was under 0.2 in both early and late S-phase, indicating that the level of overlap with these regions was less than one fifth of the average throughout the nucleus. Similar depletion was also observed during G1 and G2 phase, suggesting that the scarcity of CDC45 foci within the PCH domain isn’t simply a consequence of CDC45 sequestration at euchromatic initiation sites in early S-phase.

Notably, the HP1a-rich PCH domain has far less overlap with CDC45 foci than was measured with PCNA foci in live cells (compare Fig. 3B and Fig. 2C). While the FOC value for PCNA foci in high HP1a intensity bins is only a fraction of the peak FOC in lower intensity bins and neighboring regions, it does not consistently exhibit true depletion below the nuclear average, as would be indicated by FOC values of less than one. This is in contrast to the broad depletion of CDC45 in high HP1a bins observed at all stages of the cell cycle. While EdU labelling was primarily used to discern cell-cycle stage, it also offers an orthogonal approach to visualize elongation sites and their relationship to the PCH domain. More importantly, the amount of HP1a-overlap with CDC45 foci and replication foci (EdU) can be compared in the same cells. As expected, EdU foci had a similar distribution and morphology to PCNA foci observed in live cells, and their relationship with the PCH domain appeared qualitatively similar. Many cells in late-S phase appeared to have a “shell” of EdU foci surrounding the PCH domain, similar to what was observed for PCNA at the onset of late S-phase in live cells, while others had a morphology resembling those in the latter half of late-S, with EdU foci interspersed in channels of relatively low HP1a intensity sandwiched between higher intensity regions (Fig. 3C). EdU foci exhibited more overlap and closer proximity to high-intensity regions than was observed in live cells (Fig. 3D), but this is likely an artefact of decreased effective resolution due to cell shrinkage during fixation or processing.

CDC45 colocalization with the PCH domain was likely also affected by the shrinking, but this would be expected to bias the result towards greater colocalization with the domain. The relatively strong depletion of CDC45 foci within the PCH domain detected despite this issue strengthens the result, especially when it is compared to that observed for both EdU and PCNA. We conclude that CDC45 foci are present at reduced levels within the PCH domain throughout the cell cycle, and suggest that this may contribute to the replication dynamics observed in live cells.

## Discussion

Our results reveal that heterochromatin replication proceeds in an outside-in progression, beginning at the periphery of the PCH domain at the onset of late S-phase and later extending into low-HP1a regions interspersed throughout its interior, consistent with the progression reported in recent mammalian studies^8,9,26^. Combined with the observed depletion of CDC45 foci within the PCH domain, these results support a model in which reduced levels of CDC45 in the PCH domain limit the frequency of replication initiation in its interior. This model would help to explain multiple aspects of heterochromatin replication dynamics. First, it would explain the characteristic shell pattern that is observed at the onset of late S-Phase. If the exclusion of CDC45 suppresses initiation inside the PCH domain, initiation of heterochromatin replication would be expected to preferentially occur at the periphery of the domain, which forms an interface with the nucleosol. As replication forks move inward or pull heterochromatin out, more heterochromatin would be made accessible at the advancing interface, and this process would continue until replication is complete, consistent with the observations made here and in previous work^7,9^. Second, this model could help to explain the temporal dynamics of heterochromatin replication. If replication initiation were limited to the interface of the PCH domain, only a small fraction of heterochromatin would be accessible for replication at any given time, which would be expected to delay its overall timing. While this study focused on CDC45, it is possible that other initiation factors are similarly depleted within the PCH domain, and this would be expected to strengthen the impact on replication dynamics.

### How CDC45 depletion in the PCH domain relates to the distribution of replication foci in early and late S-phase

In late G1 and at the start of S-phase, origins licensed with the MCM2-7 helicase complex greatly outnumber the initiation factor CDC45 (Fig. 4A-B). Only a small minority of these origins initiate replication over the entirety of S-phase, with an even smaller fraction active at any given point^27,28^. Licensed origins compete for the recruitment of CDC45, which was previously shown to be rate-limiting for replication initiation in species ranging from budding yeast to mammals^15–18^. The observed depletion of CDC45 within the PCH domain is therefore likely to have a disproportionately large influence on the rate of replication initiation, resulting in a reduced frequency of initiation events within the domain relative to the euchromatic regions outside it. This difference in initiation rates provides a simple explanation for the early-S pattern of replication foci, in which replication is predominantly observed throughout euchromatic regions and largely excluded from the interior of the PCH domain.

**Figure 4.**
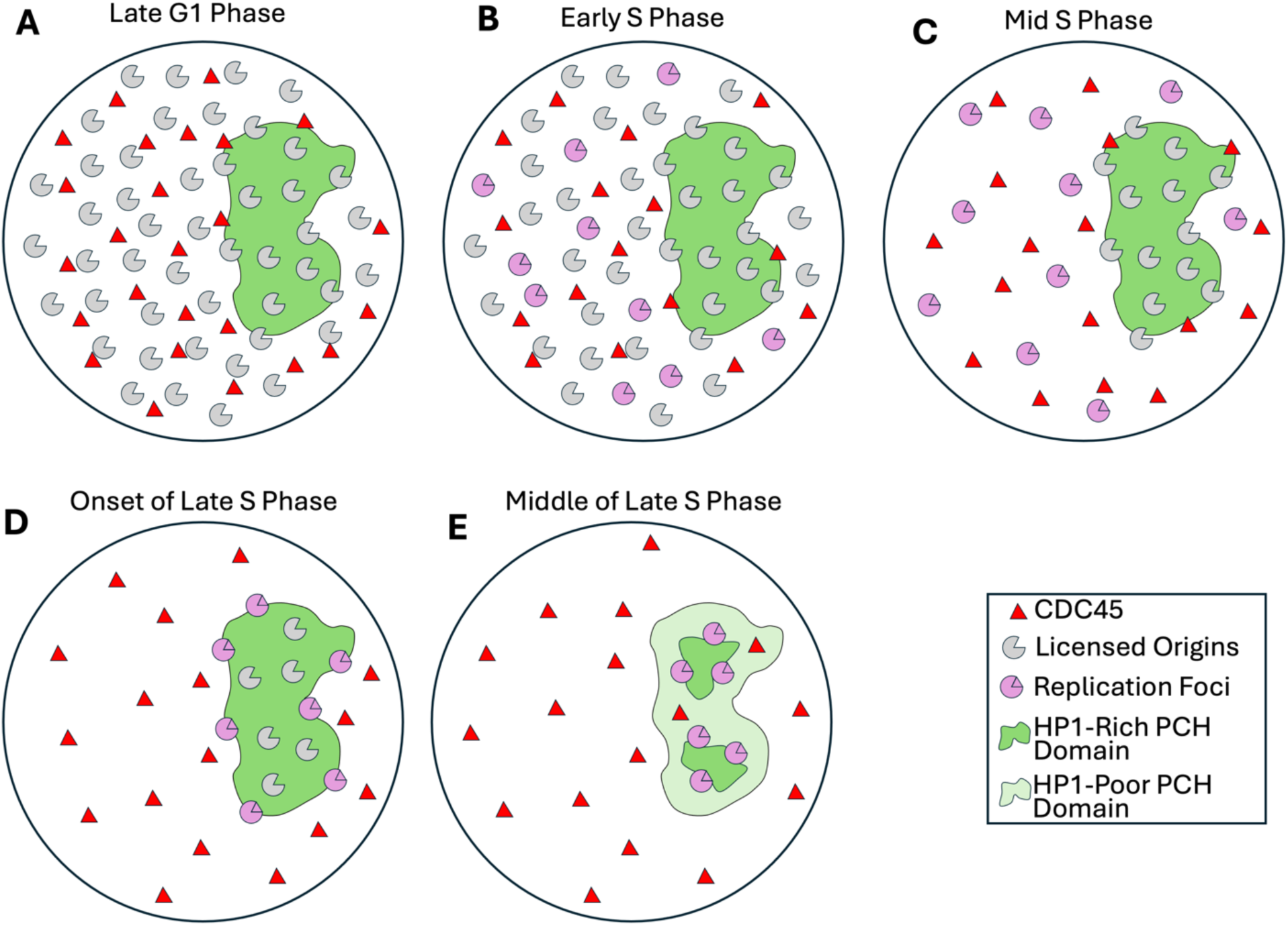
A model for the influence of CDC45 distribution on PCH replication dynamics. **A.** CDC45 foci are depleted within the PCH domain throughout the cell cycle. Licensed origins are present in excess to CDC45 throughout the nucleus, causing CDC45 to be the limiting factor for replication initiation. **B.** At the start of S-phase, the higher concentration of CDC45 foci outside the PCH domain causes euchromatic origins to fire at a higher rate than those inside the PCH domain. **C.** By mid-S Phase, the concentration of licensed origins decreases as most of the chromatin outside the PCH domain is replicated, and origin concentration becomes the limiting factor. **D.** As decreasing origin concentration limits the rate of replication initiation outside the PCH-domain, the greater concentration of origins at the interface between the PCH domain and the nucleosol leads to a higher relative rate of replication initiation there. This creates the “shell” of replication foci and marks the start of late-S and heterochromatin replication. CDC45 remains depleted within the PCH-domain, so replication is primarily initiated at the outer periphery. **E.** Replicated PCH at the periphery of the domain temporarily decondenses and/or dissociates from HP1a, allowing CDC45 to initiate replication at the periphery of unreplicated HP1a-rich PCH.

While this relationship is fairly straightforward in early S-phase, where the distribution of CDC45 is broadly similar to the distribution of replication foci, it grows more complex at the onset of late S-phase. This transition is characterized not only by a decrease in euchromatin replication, but by an increase in heterochromatin replication at the surface of the PCH domain, yet no corresponding increase in CDC45 is observed at this location. We propose that this apparent contradiction can be resolved if two effects are taken into consideration. First, as an increasingly high fraction of the euchromatin distributed throughout the nucleus is replicated during early S-phase, there is a decrease in the remaining number of licensed origins that may associate with CDC45 to initiate replication, while CDC45 is released from disassembled CMG complexes at terminated replication forks and made available for initiation at new sites^29^. Eventually, the declining number of licensed origins becomes the limiting factor for replication initiation outside of the PCH-domain. As a consequence, fewer new replication foci would form in these regions, and an increasing proportion of replication foci would be found at the domain’s surface where licensed origins remain in excess (Fig. 4C). While this effect may explain the change in the distribution of replication foci in relative terms, it is inadequate to account for the absolute increase in the amount of replication occurring at the periphery of the PCH domain at the onset of late S-phase, underlying the need for another mechanism that might be responsible for the kinetics of this this transition.

### A possible role for increasing origin firing probability in the transition to late S-phase

Past efforts to model the relationship between origin licensing, activation, and replication timing addressed a similar discrepancy by suggesting that origin firing probability increases with the progression of S-phase, keeping the overall rate of replication from declining prematurely as the number of unreplicated origins decreases^30,31^. When combined with the relative increase in the fraction of initiation events occurring at the surface of the PCH domain, this effect may explain the increased abundance of replication foci observed at this region during the transition to late S-phase. The tight coordination between the decrease in euchromatin replication and increase in PCH-associated replication foci may be particularly well-explained by a negative feedback loop that globally regulates replication initiation rates.

One intriguing candidate for such a role is the ATR-Chk1 pathway. While ATR-Chk1 is well-known for its activation of the intra-S checkpoint in response to replication stress, it is also activated at lower levels by DNA replication in unperturbed cells, where it limits excessive origin firing in early S-phase^32–34^. As licensed origins outside of the PCH domain grow sparse in mid S-phase and the rate of new initiation events begins to decline, there may be a corresponding decrease in activation of ATR-Chk1 and a relaxation of its antagonism of further origin firing, increasing the firing probability for the remaining origins. This would be expected to accelerate the completion of euchromatin replication outside the PCH domain and to further promote the relaxation of inhibition by ATR-Chk1. The further relaxation of inhibition from ATR-Chk1 would then cause an increase in the rate of origin activation at the remaining CDC45-exposed origins, which are located at the interface between the PCH domain and the rest of the nucleus.

Together, these effects would produce the abrupt transition to late S-phase and the shell-like pattern of replication foci surrounding the PCH domain observed at this time. Supporting this hypothesis, prior studies have shown that cells in early S-phase are more sensitive to the effects of ATR inhibition than those in late S-phase, and that ATR-Chk1 suppresses CDK1-mediated origin firing in early S-phase but not in late S-phase^33,35^.

### Local decondensation is consistent with internal replication patterns but insufficient to explain the initial restriction of replication to the periphery

Two distinct patterns of replication were observed during late S-phase. While replication was initially limited to the periphery of the PCH domain at the start of late S-phase, it was later observed in regions with relatively low HP1a levels interspersed between regions of higher intensity. The low HP1a intensity in these regions may reflect localized heterochromatin decondensation induced by ongoing or recent replication, partial dissociation of HP1a, or a combination of the two. While the methods used here cannot directly distinguish between these possibilities, our results closely match the pattern of locally decondensed replication sites observed within the PCH-domain in prior works, and the broader model is compatible with both potential mechanisms^8,9,26^.

Because decondensation is a known consequence of replication, it may initially seem to challenge the need for an additional mechanism to explain the relative scarcity of replication within the highly compact, HP1a-rich core of the PCH domain. Any replication occurring within these compact regions would necessarily induce local decondensation and reduce HP1a intensity, creating an apparent exclusion of replication foci from regions of high HP1a intensity. However, this is not sufficient to explain the observed distribution of foci at the start of late S-phase, when they are associated with the domain’s surface rather than decondensed regions within it. If replication initiation occurred randomly in heterochromatin and were not influenced by the organization of the PCH domain, replication foci should initially appear with comparable frequency throughout the domain, each coinciding with localized regions of low HP1a intensity caused by ongoing replication. Instead, replication foci were observed predominantly at the periphery of the domain at the onset of late S-phase, producing the characteristic shell-like pattern described above. Only later in the progression of late S-phase did replication foci appear deeper within the PCH domain at interspersed, locally decondensed regions. This sequential pattern strongly suggests that replication-induced decondensation, while influencing local chromatin structure, cannot itself explain the peripheral localization of replication foci observed at the start of late S-phase. The distribution of CDC45 offers an explanation for these dynamics, as described in the above proposed model.

### Potential mechanisms underlying the depletion of CDC45 within the PCH domain and implications for other initiation factors

While the experiments conducted are unable to determine why CDC45 is present at reduced amounts within the PCH domain, the results are consistent with the partial exclusion of CDC45. Such an effect may be mediated by the domain’s condensate properties, which have previously been shown to produce selective permeability and partially exclude inert macromolecular probes^4^. It could also be the result of affinity partitioning, in which a lack of favorable interactions with other components of the PCH domain or presence of favorable interactions with nucleoplasmic components would be expected to concentrate CDC45 in the nucleoplasm and deplete it within the PCH domain^36,37^.

The results also argue against several alternative potential explanations. Unlike PCNA, CDC45 was not observed to form foci colocalizing with sites of active replication, so its relative scarcity in high-HP1a regions of the PCH domain cannot be expected to occur solely as a consequence of replication-induced decondensation during S-phase. Another possibility is that CDC45’s localization within the nucleus might reflect its recruitment to licensed origins. If this were the case, its uneven distribution might be caused not by exclusion from the PCH domain, but by reduced recruitment to origins inside the domain or preferential recruitment to origins elsewhere. This could occur for multiple reasons. Heterochromatin-associated factors could directly or indirectly inhibit the recruitment of CDC45. For example, RIF1 is known to be enriched in the PCH domain and inhibit origin activation by antagonizing DDK activity^11^. Because DDK activity is required for CDC45 recruitment, RIF1 could also suppress CDC45 loading, although this has not yet been directly tested. Alternatively, if licensed origins are sparser in heterochromatin than in euchromatin—as has been suggested by some results but contested in others^28,38–40^—this could also reduce CDC45 recruitment in the PCH domain.

However, these mechanisms are less consistent with our results for multiple reasons. First, if CDC45’s localization directly reflected its recruitment to or association with licensed origins, its localization should shift to the PCH domain during heterochromatin replication in late S-phase, after the exhaustion of euchromatic origins outside of the PCH domain—yet no such shift was observed. Second, CDC45’s exclusion from the PCH domain was also observed in G1 cells that lacked origin activation or replication and in G2 cells with no licensed origins at all. Therefore, while differences in origin licensing and activation may contribute to replication dynamics and timing, they cannot explain the reduced levels of CDC45 within the PCH domain and their persistence throughout the cell cycle.

If selective permeability, affinity partitioning, or a similar mechanism is responsible for CDC45’s partial exclusion from the PCH domain, it’s possible and perhaps likely that other factors required for replication initiation are also affected by these mechanisms and distributed in a similar manner. If multiple initiation factors are partially excluded from the PCH domain, this would be expected to produce a synergistic effect on the rate of assembly of the mature CMG complex and consequent origin firing. This could help to explain how the partial exclusion of CDC45 would result in a near total absence of replication foci within the PCH domain until the final stage of late S-phase, following extensive replication and decondensation at the domain’s periphery. If licensed origins are more frequent in euchromatin than heterochromatin, this might also contribute to the observed dynamics in a similar fashion, although the stoichiometric excess of licensed origins makes it likely that its influence on replication timing would be lesser than that of CDC45 and other initiation factors.

### Interpretation of CDC45 foci and their relationship to replication initiation and elongation

The nuclear localization of CDC45 is in broad agreement with past results that have visualized the protein, but its distribution within the nucleus has been more variable between studies with differing cell types and visualization techniques. Some studies observed colocalization with sites of ongoing replication, as might be expected based upon CDC45’s membership in the replicative CMG complex, but the studies reporting this localization did so only after removal of the soluble protein fraction prior to fixation, within the atypical context of Chorion amplification, or after potentially disruptive synchronization with thymidine blocks^41–43^. Other studies are in better agreement with the localization observed in the present work, finding broad nuclear localization with diffuse or punctate distribution that does not colocalize with replication foci^16,18,44^. The pan-nuclear distribution seems to be more common when CDC45 is visualized without removal of the soluble protein fraction, as is the case in the present study. A lack of visible association with replication foci despite an essential biochemical role in replication would not be unique to CDC45, as it has been well documented for other replication proteins, most notably in the case of the MCM complex^18,45,46^. This paradoxical lack of colocalization may be caused by epitope masking within the fully assembled CMG complex, although this explanation is made less likely in the present study by prior validation of the antibody used here through immunoprecipitation of the full CMG complex^43,47^. It might also result from a large fraction of CDC45 that is soluble, bound to origins that have not yet fired, or associated with other structures, in which case the signal from these other populations might drown out the signal that is associated with replication foci. This possibility is supported by the past findings that only a small fraction of CDC45 is stably bound to chromatin at any given time and that CDC45 frequently binds to licensed origins for lengthy periods without firing^41,48^.

Whatever the cause, the lack of colocalization with replication foci was an advantage in the context of the current work, where CDC45 was selected because of its rate-limiting role in replication initiation rather than its association with replication foci. If CDC45 had strongly colocalized with replication foci, it would have simply recapitulated the results produced with PCNA and would not have been informative.

The observed CDC45 foci are also unlikely to be specific to replication initiation sites, as they were present throughout the cell cycle independent of replication or licensing. Segmentation of CDC45 foci was performed not to identify putative initiation sites but rather to reduce non-specific background signal, facilitate analysis with FOC-Map, and ensure methodological consistency with the approach used for replication foci to allow direct comparison of results. Our model uses the localization of CDC45 foci primarily as a tractable proxy for assessing relative initiation potential rather than direct initiation events. Follow-up studies should seek to corroborate the localization observed here using orthogonal methods to visualize CDC45, ideally with endogenous labelling in live cells. No prior work has directly assessed the relationship between CDC45 and the PCH domain, although the figures in at least one prior work appear to display relatively little CDC45 within the DAPI-bright regions characteristic of PCH domains^18^.

### The role of HP1a, outstanding questions, and future directions

Although our experimental design does not enable any conclusions about HP1a’s possible contribution to the observed dynamics, several pieces of evidence make it a promising candidate for further study of the spatio-temporal regulation of PCH replication. HP1a’s central role in the organization and establishment of the PCH domain through condensate formation suggest that it may help to orchestrate the domain’s exclusion of CDC45, and possibly of other replication factors whose localization with respect to the PCH domain have not yet been characterized. If HP1a is at least partially responsible for CDC45’s exclusion, this could provide a mechanism for its documented modulation of PCH replication timing^6,49^. It would be informative to test whether depletion of HP1a lessens or abolishes CDC45’s exclusion, particularly in comparison to perturbations that reduce the compaction of PCH without disrupting its recruitment of HP1a. Future work should also determine whether low levels of CDC45 are more highly correlated with HP1a than they are with chromatin compaction. It would also be of interest to assess whether artificially increasing CDC45 levels within the PCH domain would be sufficient to cause early replication of PCH, or an interspersed pattern of replication within the domain at times when replication would ordinarily be restricted to the periphery. Such a perturbation can be achieved by tagging CDC45 with one of the recently identified weakly binding variants of the HP1a interacting motif that could facilitate access to the domain without causing strong enrichment^12^.

This work provides additional support for an emerging consensus that heterochromatin replication occurs not only at the outer periphery of the PCH domain, but also in locally decondensed internal regions. It additionally suggests that large scale relocalization of internal sequences to the surface of the domain may not be required for replication. At the same time, our study validates earlier observations of replication foci strongly enriched at the periphery of the PCH domain and is one of only two studies that situates this pattern within a consistent temporal sequence using time-lapse imaging in live cells. It also extends this understanding by suggesting a possible mechanistic framework that relates these spatio-temporal dynamics to the newly observed exclusion of CDC45 from the PCH domain. Future work should seek to validate these findings and explore the new questions they raise, particularly regarding HP1a’s role in the exclusion of CDC45 from the PCH domain and the distribution of other replication initiation factors.

## Methods

### Transient transfection of S2R+ Drosophila cells

Cultured S2R+ cells (DGRC Stock 150; RRID:CVCL_Z831) in log-phase growth were resuspended in Schneider’s Medium (Gibco Cat. 21720024) supplemented with 10% Fetal Bovine Serum (FBS) at a concentration of 1×10^6^ cells/mL. 8-well chamber slides (Ibidi Cat. 80826) were seeded with 200 uL of resuspended cells per chamber, and cells were allowed to recover and adhere overnight at 25°C. Cells were transfected the following day with plasmids containing mScarletI-PCNA and mEGFP-HP1a under the control of the Copia promoter (pCopia). Plasmid transfection was performed with TransIT-2020 (Mirus Cat. MIR 5400) transfection reagent using a modified version of the manufacturer’s protocol. Briefly, each well was transfected with 100 ng of each plasmid (200 ng total) complexed with 0.2 μL of TransIt-2020 transfection reagent in 40 uL of Serum-Free Schneider’s Medium. Transfection mixture was added dropwise after incubating at room temperature for 30 minutes.

### Time-lapse image acquisition and processing

Time-lapse imaging was performed 48-72 h after transfection. Images were collected on a Zeiss LSM880 with a Plan-Apochromat 63×/1.4 NA Oil DIC M27 objective and an Airyscan Detector in Fast Mode. Cells in early S-phase with numerous dispersed PCNA foci were selected for imaging to allow subsequent progression through late S-phase to be captured. Z-stacks were collected every 30 mins for 12-16 h with 2x line averaging and frame-sequential channel-switching. A zoom of 4.0 was used with the SR “optimal” presets for frame size and Z-interval, giving voxel dimensions of 43 x 43 x 211 nm. Both channels were imaged with a pixel dwell time of 0.55 μs and Master Gain of 775 with the “MBS 488/561” main beam splitter. mScarletI-PCNA was excited with the 561 nm laser line at 0.7% power with the “BP 570-620 + LP 645” emission filter. mEGFP-HP1a was excited with the 488 nm laser line at 0.5% power with the “BP 420-480 + BP 495-550” emission filter. Images were deconvolved by 3D Airyscan Processing in Zen Black (Zen 2.3 SP1) using the default processing strength.

Segmentation of nuclei was done in Arivis Vision4D (version 3.6) by setting a threshold for diffuse nuclear HP1a, which is far above the cytoplasmic and extracellular background. Prior to segmentation, the HP1a channel was denoised with a discrete Gaussian filter with a diameter of 4.95 μm. Segmentation was then performed with the Watershed method, using a threshold of 150, a diameter of 8.75 μm, and a split sensitivity of 0.5%. Segments were then tracked to group the segments for each nucleus across timepoints. The segments for each nuclear track were then converted to saturated masks and exported as .ome.tiff files.

Nuclei were then normalized within a custom FIJI macro to reduce the effects of photobleaching and variability in expression levels between transfected cells. For the mEGFP-HP1a images, nuclear intensity values were divided by the average nuclear intensity at each timepoint, yielding the relative intensity of each nuclear voxel. For the mScarletI-PCNA channel, a single rescaling factor was calculated from the average mean nuclear intensity of the first three timepoints and applied uniformly to all timepoints. This approach was chosen to correct for differences in expression level between cells while preserving changes in overall PCNA intensity. Following normalization, segmentation of PCNA foci from t was performed in Arivis Vision4D using an adaptive mean threshold with a diameter of 0.341 μm, a local threshold of 0.25, and 55% thinning.

To enable comparison between cells with different rates of S-phase progression, the time-lapse data from each cell were temporally aligned by expressing each timepoint as a percentage of the duration of late S-phase. The onset of late S-phase was identified by visual inspection as the timepoint at which PCNA foci converged at the PCH domain, and its completion was defined as the last timepoint at which segmented PCNA foci were present. Timepoints in early and mid S-phase precede the onset of late S-phase and therefore have negative progression values. From each time-lapse, the timepoints closest to -20%, 20%, 60%, and 100% progression through late S-phase were selected for subsequent analysis with FOC-Map.

### Derivation and Definition of the Fractional Overlap Coefficient

The FOC quantifies the spatial association between segmented objects in one channel and bins in the other by comparing observed overlap to that expected under a null model of random distribution. To determine how segments in one channel (e.g. replication foci, referred to here as the red channel) are distributed across bins in the other (e.g. HP1a intensity or distance bins, referred to as the green channel), one could calculate the fraction of red segment volume that overlaps each green bin:

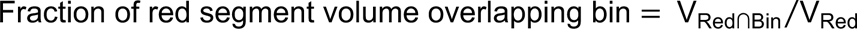

where V_Red_ is the total volume of segments in the red channel and V_Red∩Bin_ is the volume of overlap between the red segments and the bin. However, when bins differ in volume, larger bins will on average harbor a proportionally larger fraction of the red segments even if the red segments are randomly distributed, preventing meaningful comparison between bins of different sizes. To account for this, the FOC normalizes the observed fractional overlap by the fraction of the total region of interest (ROI) volume occupied by the bin:

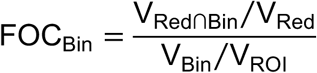

where V_Bin_ is the volume of the bin and V_ROI_ is the volume of the region of interest (ROI), defined as the volume in which two signals could potentially co-occur (e.g. the nucleus, for two nuclear proteins). The FOC can also be understood through the inverse formulation: the fraction of the bin volume overlapping red segments, normalized by the fraction of the ROI volume occupied by red segments:

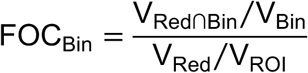

Both of these approaches are mathematically identical, and can be simplified to give the general equation below:

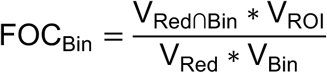

### Live-cell FOC-Map analysis

FOC-Map was performed to quantify the distribution of replication foci with respect to the HP1a-rich PCH domain. FOC-Map processing was implemented in Matlab (Mathworks R2023b) using custom scripts. The normalized HP1a intensity images and binary PCNA foci masks produced by the preceding steps were used as inputs. Within each nucleus, voxels with HP1a intensity above the nuclear background were sorted into 1000 percentile intensity bins (0.1% increments from the 0th to 99.9th percentile), and the average HP1a intensity of voxels in each bin was recorded. Voxel coordinates were converted from image indices to nanometers using the Airyscan-processed voxel dimensions (42 × 42 × 211 nm) to account for anisotropic voxel geometry. For each voxel in a given percentile intensity bin, all nuclear voxels were binned by their distance from that voxel in 42 nm intervals. Within each distance bin, the total number of nuclear voxels and the number of segmented PCNA foci voxels were counted. This process was repeated for every voxel in the intensity bin, and the resulting counts were pooled across voxels within each distance bin. The procedure was then repeated for each percentile intensity bin. To generate pooled results for each timepoint, the nuclear voxel counts and PCNA foci voxel counts for each combination of intensity bin and distance bin were summed across all nuclei at that timepoint, as were the total PCNA foci and nuclear volumes. The FOC was then calculated from these pooled counts for each combination of intensity bin and distance bin. The average HP1a intensity for each percentile bin was recalculated from the pooled intensity values across all contributing nuclei.

For Figure 2B, the FOC values at zero distance (direct overlap) were extracted for each intensity bin and plotted against the corresponding average HP1a intensity, expressed as fold nuclear average. For Figure 2C, the full matrix of FOC values across all intensity and distance bins was displayed as a pseudocolor heatmap with HP1a intensity on the X-axis (scaled by average bin intensity rather than percentile rank), distance on the Y-axis, and FOC encoded as color. Percentile bins left empty due to limited bit-depth in the intensity values were excluded from the heatmap prior to plotting. Heatmaps were plotted using interpolated shading with a color scale range of 0 to 25. Analysis code used in this study is available at https://github.com/collin-hickmann/FOC-Map.

### EdU-labelling and immunofluorescence

Clean and sterile coverslips (22×22mm 1.5H) were incubated in 100% Fetal Bovine Serum (FBS) for 2 h at 37°C to facilitate adsorption of extracellular matrix components and promote cell adherence. After rinsing with 1x PBS, coverslips were transferred to 6-well plates and seeded with 2 mL of 1×10^6^ S2R+ cells/mL in Schneider’s Medium supplemented with 10% FBS. Cells were allowed to adhere overnight at 25°C. The following day, cells were treated with 10 μM EdU in conditioned medium for five minutes, fixed for five minutes in 4% paraformaldehyde (PFA) in 1x PBS, then permeabilized and washed in 1xPBS + 0.4% TritonX-100 (PBST) three times for five minutes. Residual fixation was quenched by incubation in blocking buffer (1x PBST + 5% FBS) for one hour at room temp. Labelling with Alexa647 picolyl azide was performed with the Click-iT Plus kit (Invitrogen Cat. C10640) according to the manufacturer’s protocol, but the Click-labeling step was performed twice to increase the signal and compensate for the brief duration of pulse-labeling. After labeling, fixed coverslips were stored overnight in blocking buffer at 4°C.

Coverslips were then incubated at 4°C with blocking buffer containing 1 μg/mL monoclonal mouse anti-HP1a (C1A9 from DGRC) and a 1:400 dilution of affinity purified rabbit anti-CDC45, which was kindly provided by Mike Botchan and has been validated to recognize CDC45 within the fully assembled CMG complex ^47^. Primary antibody incubation was continued overnight at 4°C followed by 3x 5 min washes in room temperature PBST. Secondary antibody staining was done with 1:500 dilutions of Alexa 568 goat anti-mouse (2 mg/mL; Invitrogen Cat. A21202) and Alexa488 goat anti-rabbit (2 mg/mL; Invitrogen Cat. A11034) in blocking buffer incubated at room temperature for 90 minutes. Coverslips were again washed 3x 5 minutes in PBST, then post-fixed in 4% PFA in 1x PBST for 5 minutes. PFA was then washed off with 3x 5 min incubations in PBST and quenched in blocking buffer for 1 hour. Nuclei were counter-stained with DAPI in 1x PBS for five minutes, quickly rinsed in 1xPBS, and mounted on slides with SlowFade Glass (Thermo Cat. S36917). Mounted slides were sealed with nail polish and stored at -20°C before imaging.

### Fixed cell image acquisition and processing

Z-stacks were acquired on the Zeiss LSM880 with a Plan-Apochromat 63×/1.4 NA Oil DIC M27 objective and an Airyscan Detector in Superresolution mode. Images were acquired using 2x line averaging and frame-sequential channel-switching with a pixel dwell time of 1.04 μs, a zoom of 7.5, and the optimal frame size in SR mode, giving voxel dimensions of 35 x 35 x 159 nm. All channels were imaged with a master gain of 800 and bit-depth of 16 with the MBS 488/561/633 and MBS-405 main beam splitters. The SBS LP 660 secondary beam splitter was used only when imaging Alexa647-EdU to prevent bleed-through from Alexa568. Alexa647-EdU was excited with the 633 nm laser at 2% power with the BP 570-670 + LP 645 emission filter. Alexa568-labeled anti-HP1a was excited with the 561 nm laser line at 0.3% power using the BP 420-480 + BP 495-620 emission filter. Alexa488-labelled anti-CDC45 was excited with the 488 nm laser line at 0.5% power. DAPI was excited with the 405 nm laser at 0.2% power and the BP 420-480 + BP 495-550 emission filter.

Images were deconvolved by 3D Airyscan Processing (SR Mode) using processing strengths of 7.3 for Alexa647-EdU, 7.5 for Alexa568-HP1a, 7.0 for Alexa488-CDC45, and 7.0 for DAPI in Zen 2.3 Blue Edition. Chromatic aberration was measured from images of a tetraspeck bead slide acquired with the same settings used for fixed cells. After airyscan processing, the shifts between channels were by corrected by lateral channel alignment in Zen Black using the average shifts measured from three bead images. After correction of chromatic aberration, nuclear segmentation was performed in Arivis Vision4D. The HP1a and DAPI channels were each smoothed with a discrete Gaussian filter (4 μm diameter), summed, and the resulting combined image was segmented using the Otsu method.

Background correction and normalization of fixed-cell images were performed with a custom Fiji macro. A constant background intensity was estimated independently for each channel using a conservative approach based on the minimum local average intensity in the image. The top five and bottom twenty z-slices were excluded to avoid edge artifacts from Airyscan processing and elevated background near the coverslip surface. A mean filter was applied in XY (radius of 20 pixels) and in Z (radius of 2 slices), and the resulting smoothed stack was reduced by minimum-intensity z-projection. The minimum value in the projection was taken as the estimated background for each channel and subtracted as a constant from all voxels, with any resulting negative values set to zero. The HP1a channel was normalized by dividing nuclear voxel intensities by the mean nuclear intensity, yielding relative intensity values as described for the live-cell images. To enable consistent segmentation of foci across cells with varying absolute intensities, the EdU and CDC45 channels were linearly rescaled within each nucleus such that the maximum nuclear voxel intensity was set to a fixed standard of 10,000. Segmentation of EdU and CDC45 foci was then performed in Arivis Vision4D using the adaptive mean method with a diameter of 0.213 μm, a local threshold of 750, and 55% thinning. The resulting segments were converted to saturated masks and exported as OME-TIFF files.

### Classification of cell cycle phase in fixed cells

Cells in S-phase were distinguished from G-phase cells by the presence of EdU foci. Cells in each image were manually annotated as being in early or late S-phase using roughly the same strategy as in live cells, where cells with numerous dispersed replication foci were annotated as being in early or mid S-phase, while those with a smaller number of replication foci near the PCH domain were classified as being in late S-phase. Cell cycle classification was checked against the count of segmented replication foci and integrated DAPI signal, which both correlated well with S-phase progression, with manual staging being used to more accurately classify the few outliers where cell cycle stage was not captured by these more objective methods.

G-phase cells were classified as G1 or G2 based on integrated nuclear DAPI intensity, which serves as a proxy for total DNA content. A histogram of integrated DAPI among G-phase cells revealed a clear bimodal distribution, with one peak between 50 and 59 million arbitrary units and a second peak between 100 and 119 million arbitrary units, consistent with pre- and post-replication DNA content respectively. Of the 228 cells for that were classified for cell cycle stage, 4 EdU-negative (G-phase) were excluded from analysis because their DNA content fell between the two peaks and did not permit a reliable classification as G1 or G2.

### FOC-Map analysis of fixed cells

FOC-Map analysis of fixed-cell images was performed using the same approach described for live-cell time-lapse data, with modifications to account for differences in image acquisition parameters and experimental design. The analysis was implemented in custom MATLAB scripts and followed the same computational workflow of percentile intensity binning, distance binning, and calculation of the Fractional Overlap Coefficient (FOC). HP1a intensity within each nucleus was normalized to the mean nuclear value, as was done for live cells. Percentile intensity bins of 0.05% were used (compared to 0.1% for live cells), and distance bins were set to 35.42 nm, matching the lateral voxel dimensions of the fixed-cell Airyscan images. The analysis was performed independently for EdU foci and CDC45 foci, each measured against the HP1a channel, using the binary foci masks generated during segmentation.

Rather than aligning cells by temporal progression through late S-phase as was done for live-cell time-lapse data, fixed cells were grouped by cell-cycle stage and pooled within each group. Pooling was performed by summing the raw voxel counts for each combination of intensity and distance bin across all cells in a given stage prior to calculating the FOC, following the same pooling strategy used for live cells. The resulting FOC-Maps represent the aggregate spatial relationship between foci and the HP1a-labeled PCH domain for each cell-cycle stage.

## Acknowledgements

Sincere thanks to all members of the Karpen lab for feedback and guidance. Thanks to Mike Botchan for sharing antibodies against CDC45 and other replication factors, and to Serafin Colmenares for the plasmids used in transient transfections. This work was supported by funding from the National Institutes of Health (grant numbers R01 GM117420, T32 HG000047, T32 GM132022 and R35 GM139653).

## Supplemental Material

**Supplemental Figure 1.**
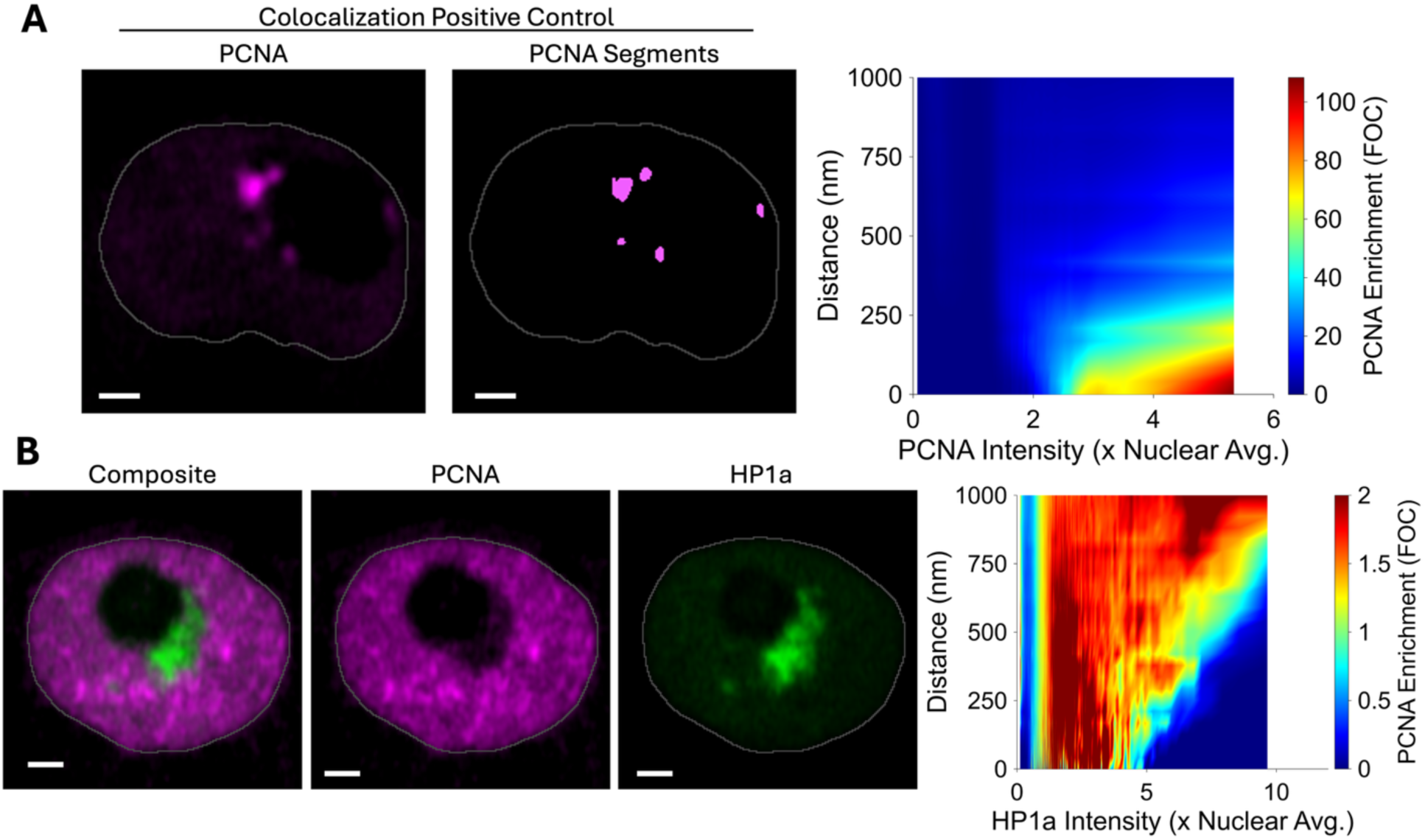
FOC-Map visualization of colocalization and exclusion. **A.** FOC-Map analysis of a positive control for colocalization visualizing the normalized overlap (FOC) between segmented PCNA foci and PCNA percentile intensity bins. **B.** A live S2R+ cell in early S-phase, when PCNA foci are enriched in euchromatic regions of the nucleus and depleted within the PCH domain. The FOC-Map showing the distribution of PCNA Foci over bins of varying HP1a intensity (x-axis) and bins of varying distance to each intensity bin (y-axis) for this cell. An FOC well below 1 indicates depletion (dark blue) in the high intensity bins (5-10x Nuc. Avg.), while the FOC above 1 reveals enrichment (orange-red) of PCNA foci in bins of low-moderate HP1a intensity (0-5x Nuc. Avg.).

## Notes

https://github.com/collin-hickmann/FOC-Map

### Competing Interest Statement

The authors have declared no competing interest.

